# The bovine alveolar macrophage DNA methylome is resilient to infection with *Mycobacterium bovis*

**DOI:** 10.1101/238816

**Authors:** Alan Mark O’Doherty, Kevin Rue-Albrecht, David Andrew Magee, Simone Ahting, Rachelle Elizabeth Irwin, Thomas Johnathan Hall, John Arthur Browne, Nicolas Claude Nalpas, Colum Patrick Walsh, Stephen Vincent Gordon, Marcin Wlodzimierz Wojewodzic, David Evan MacHugh

## Abstract

DNA methylation is pivotal in orchestrating gene expression patterns in various mammalian biological processes. Perturbation of the bovine alveolar macrophage (bAM) transcriptome, due to *Mycobacterium bovis* (*M. bovis*) infection, has been well documented; however, the impact of this intracellular pathogen on the bAM epigenome has not been determined. Here, whole genome bisulfite sequencing (WGBS) was used to assess the effect of *M. bovis* infection on the bAM DNA methylome. The methylomes of bAM infected with *M. bovis* were compared to those of non-infected bAM 24 hours post-infection (hpi). No differences in DNA methylation (CpG or non-CpG) were observed. Analysis of DNA methylation at proximal promoter regions uncovered >250 genes harbouring intermediately methylated (IM) promoters (average methylation of 33–66%). Gene ontology analysis, focusing on genes with low, intermediate or highly methylated promoters, revealed that genes with IM promoters were enriched for immune-related GO categories; this enrichment was not observed for genes in the high or low methylation groups. Targeted analysis of genes in the IM category confirmed the WGBS observation. This study is the first in cattle examining genome-wide DNA methylation at single nucleotide resolution in an important bovine cellular host-pathogen interaction model, providing evidence for IM promoter methylation in bAM.

## Introduction

Infection with *Mycobacterium bovis*, the causative agent of bovine tuberculosis (BTB), accounts annually for more than $3 billion of losses to global agriculture through lost productivity and disease control costs ^1^. There is also evidence suggesting that the burden of *M. bovis* as the cause of zoonotic tuberculosis in humans may be underestimated ^2^, which highlights the need for a more detailed understanding of the impact of *M. bovis* in both cattle and humans. Unravelling host cellular processes that are perturbed or manipulated by intracellular pathogens is an important area of research in infection biology, particularly for disease control and the development of next-generation diagnostics and prognostics. In this regard, host cell epigenetic modifications induced, either as a component of the response to *M. bovis* infection, or as an immunoevasion strategy by the pathogen itself, remain to be fully elucidated ^3^.

Modifications to the genome, such as DNA methylation and histone tail modifications, in combination with RNA-mediated regulatory mechanisms are fundamental in modulating tissue-specific gene expression ^4–6^. Epigenetic gene regulation represents an important framework for understanding how environmental stimuli are disseminated to the transcriptome and preserved through subsequent somatic cell divisions ^5^. DNA methylation (5-methylcytosine), the most widely studied genome modification, is involved in a variety of cellular processes including genomic imprinting, X-chromosome inactivation, chromosome stability and gene transcription ^7^ and has been proposed to be influenced by external stimuli across a wide range of biological contexts ^8–12^. Therefore, we hypothesised that changes to DNA methylation may be involved in the bovine host response to infection with *M. bovis*; this mechanism has previously been proposed for human tuberculosis caused by infection with *Mycobacterium tuberculosis* ^13^.

Host epigenomic plasticity to *M. tuberculosis* has been reported previously^14,15^. Sharma and colleagues showed that non-CpG loci in the host genome were hypermethylated following reduced representation bisulfite sequencing (RRBS) analysis of THP-1 macrophages (a human monocytic cell line) infected with *M tuberculosis* ^14^. In addition, Zheng *et al*. ^15^ demonstrated that interleukin gene promoter sequences, and their receptors, were associated with hypermethylation following analysis of THP-1 cells infected with clinical strains of *M. tuberculosis*, using the human inflammatory response methyl-profiler DNA methylation PCR array. Furthermore, it has been demonstrated that DNA methylation is associated with hypoxic survival of *M. tuberculosis* ^16^. Most recently, Doherty *et al*. reported that there were in excess of 750 differentially methylated regions between *M. bovis*-infected and healthy cattle in a study using RRBS to examine CD4^+^ T lymphocytes isolated from circulating blood samples ^17^.

A range of studies have highlighted the impact of infecting microorganisms on host DNA methylation patterns. For example, distinct DNA methylation changes have been observed in macrophages infected with the intracellular protozoan *Leishmania donovani*, the causative agent of visceral leishmaniasis ^18^. In addition, global DNA methylation changes have been detected in human neutrophils infected with *Anaplasma phagocytophilum*, which causes granulocytic anaplasmosis ^19^. Finally, it has been proposed that, during chronic *Helicobacter pylori* infection in humans, functional *H. pylori* DNA methyltransferases enter host epithelial cells and methylate their recognition sequences in chromosomal DNA, potentially contributing to the pathogenesis of gastric adenocarcinoma or lymphoma of the mucosa-associated lymphoid tissue ^20^.

Our group has previously revealed the impact of *M. bovis* infection on the mammalian alveolar macrophage gene expression, demonstrating that the bAM transcriptome is substantially reprogrammed as a consequence of both host-driven defence responses and mycobacterial-induced perturbation and manipulation of cellular processes ^21–24^. However, the effect of *M. bovis* on the bovine host epigenome, specifically the DNA methylome of bAM, remains unexplored. Recent work has shown that intracellular microbial infection can lead to alterations of the host DNA methylome; therefore, for the present study we used WGBS to test the hypothesis that bAM DNA methylation patterns are altered during the earliest stage of *M. bovis* infection in cattle.

## Materials and Methods

### Ethics statement

All animal procedures were performed according to the provisions of the Cruelty to Animals Act of 1876 and EU Directive 2010/63/EU. Ethical approval was obtained from the University College Dublin Animal Ethics Committee (protocol number AREC-13–14-Gordon).

### Isolation and infection of bovine alveolar macrophages

Isolation and purification of bAM from cattle was performed as previously described by our group ^21,23^ and is summarized in Fig. 1. Briefly, total lung cells were harvested by pulmonary lung lavage with Hank’s Balanced Salt Solution (Invitrogen, Life Technologies) following the removal of lungs from eight unrelated Holstein-Friesen male calves. Total lung cells were washed and cultured for 24 h at 37 °C in R10^+^ media (RPMI 1640 medium supplemented with antibiotics [Invitrogen]). After incubation, cells were prepared for infection by dissociation and seeding at 5 × 10^5^ viable cells/well, for each biological replicate. The purity of the seeded macrophages was confirmed by flow cytometry using anti-CD14 antibody. bAM were infected with *M.bovis* strain AF2122/97 at a multiplicity of infection (MOI) of 10 bacilli per alveolar macrophage as described in detail previously ^21,23^. These previous studies used comparative RNA-seq-based transcriptomics and targeted quantitative assays (RT-qPCR and multiplex ELISA) of several NF-κB-inducible pro- and anti-inflammatory cytokines and chemokines, including CCL-4, IL-1β, IL-6, IL-10 and IL-12, to verify that at 24 hours post-infection (hpi) *M. bovis*-treated bAM cells were infected and had internalised bacilli ^21,23^.

**Figure 1.**
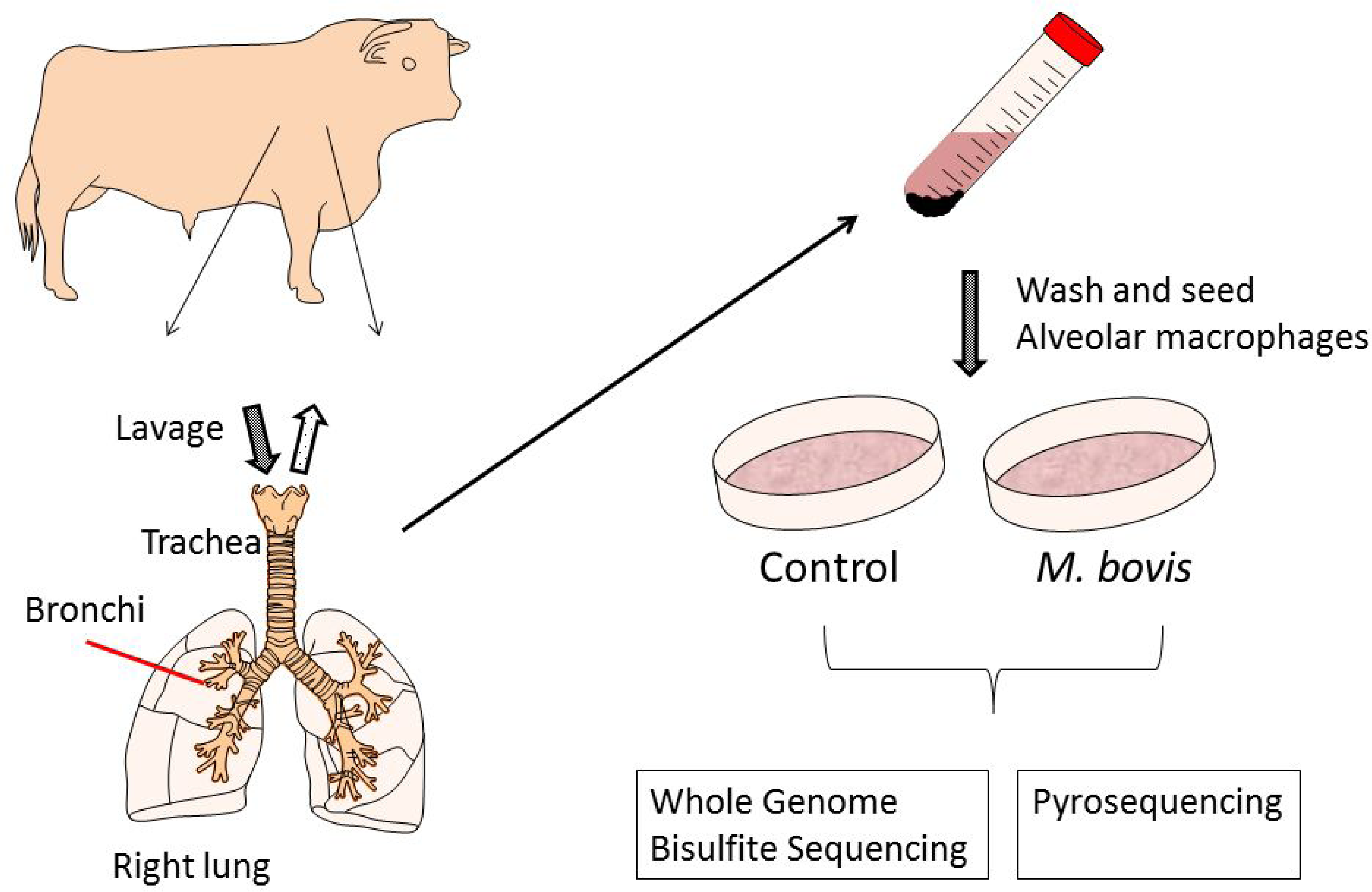
Schematic representation of sample preparation. Bovine alveolar macrophages (bAM) were isolated, post-mortem, from the lungs of age-matched male Holstein-Friesian calves by lavage. Purity of the cells was confirmed using flow cytometry with anti-CD14. Isolated cells were washed and seeded for 24 h prior to infection. Infected bAM were exposed to *M. bovis* at a multiplicity of infection ratio of 10:1 for 2 h. After 2 h the media was replaced in control and infected samples and cells were harvested after 24 h for analysis of DNA methylation. This figure was prepared by A.M.O’D. using the Biomedical PPT toolkit suite (www.motifolio.com).

### Isolation of DNA and library preparation

DNA was extracted from *M. bovis*-infected bAM 24 hpi (*n* = 8) and from control bAM (*n* = 8) at the same time point using the DNeasy kit (Qiagen) according to the manufacturer’s recommendations. DNA was quantified using the Qubit dsDNA HS Assay Kit (ThermoFisher Scientific). Libraries were prepared for WGBS using the post–bisulfite conversion library preparation method for methylation analysis (EpiGnome™ Methyl-Seq Kit, Epicentre, Illumina) according to the manufacturer’s instructions. Genomic DNA (50 ng) isolated from *M. bovis*-infected or non-infected bAM (24 hpi) was bisulfite-modified (EZ Methylation-Direct Kit, Zymo) according to the manufacturer’s guidelines. DNA synthesis was performed by mixing bisulfite-converted DNA with 2 μl DNA synthesis primer, incubating at 95 °C for 5 min, cooling on ice, followed by addition of 4 μl EpiGnome DNA Synthesis Premix 0.5 μl, 100 mM DTT and 0.5 μl EpiGnome Polymerase. Reactions were incubated at 25 °C for 5 min followed by 42 °C for 30 min, then cooled to 37 °C for 2 min before addition of 1 μl Exonuclease I to each reaction. Following this, reaction mixtures were incubated at 37 °C for 10 min, 95 °C for 3 min and then held at 25 °C. DNA was di-tagged by adding 7.5 μl EpiGnome TT Premix and 0.5 μl DNA polymerase to each reaction and incubating at 25 °C for 30 min, 95 °C for 3 min and cooling to 4 °C. Tagged DNA was purified using the using the AMPure XP (1.6× beads, 40 μl) system. A PCR step was performed to generate the second strand of DNA, complete the addition of the Illumina adaptor sequences and incorporate an index sequence. 22.5 μl of di-tagged DNA was mixed with 25 μl FailSafe PCR PreMix E, 1 μl EpiGnome Forward PCR Primer, 1 μl EpiGnome Index PCR Primer and 0.5 μl FailSafe PCR Enzyme (1.25 U) and subjected to an initial denaturation of ds DNA at 95 °C for 1 min followed by 10 cycles of 95 °C for 30 sec, 55 °C for 30 sec and 68 °C for 3 min. Following PCR, the reactions were incubated at 68 °C for 7 min. EpiGnome libraries were purified using the AMPure (1× beads, 50 μl) system to remove primer dimers. Libraries were quantified by Qubit using the Qubit dsDNA HS Assay Kit (ThermoFisher Scientific) and library quality was assessed on an Agilent BioAnalyzer using the High sensitivity DNA assay kit (Agilent Technologies).

### Pyrosequencing

Genomic DNA was extracted from *M. bovis*-infected and control bAM (isolated from a parallel set of four animals to those used for WGBS) and quantified with the High-Sensitivity DNA Assay Kit (Agilent Technologies). DNA (200 ng) was bisulfite-modified using the EZ Methylation-Direct Kit (Zymo) and eluted in 50 μl elution buffer. Bisulfite PCR reactions were performed in 25 μl consisting of 0.2 μm each primer, 2 mM MgCl_2_, 1× PCR buffer (minus magnesium), 0.2 mM dNTPs, Platinum *Taq* DNA polymerase (Invitrogen), and 3 μl bisulfite-modified DNA. Primer sequences are detailed in Table 1. PCR cycling conditions were as follows: 95 °C for 5 min followed by 40 cycles of 30 sec each at 95 °C; either 55 °C (*TNF, NFKB2* and *IL12A*) 56 °C (*DTX4, C1QB* and *NOS2*) or 58 °C (*TLR2*) for 30 sec; 72 °C for 30 sec, and a final elongation step of 5 min at 72 °C. PCR products were verified by electrophoresis on a 2% w/v agarose gel before pyrosequencing (Pyromark Q24, Qiagen). Pyrosequencing assays were designed in-house and carried out as previously described ^25,26^. Only pyrosequencing reactions that passed Pyromark Q24 internal controls for bisulfite modification were included in the analysis. Two-tailed paired sample *t*-tests were used to assess statistically significant DNA methylation between control and *M. bovis*-infected samples.

**Table 1.**
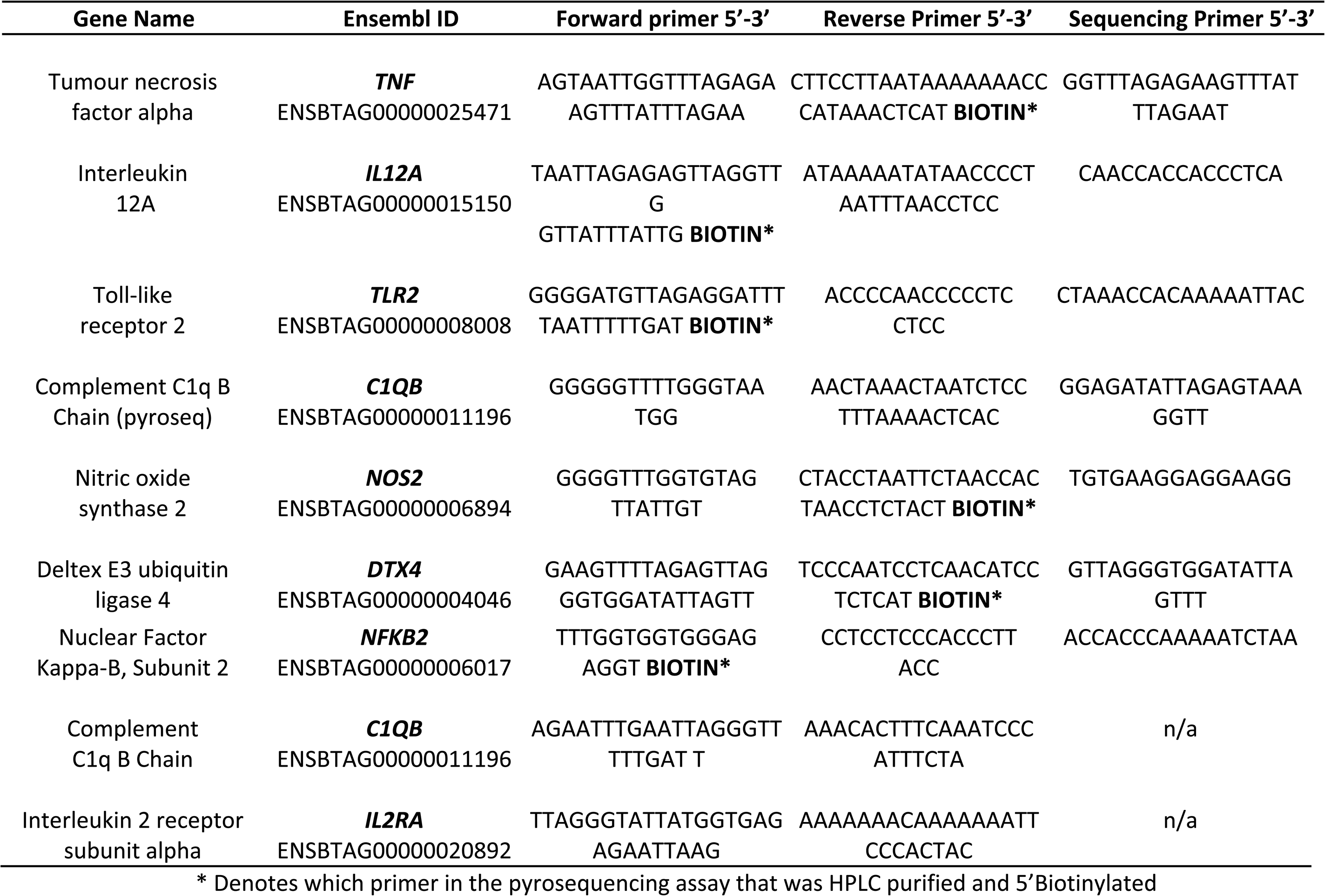
Primers use for targeted DNA methylation analysis.

### Bisulfite PCR, cloning, sequencing and combined bisulfite restriction analysis

Bisulfite-converted DNA from control and infected bAM was amplified in 25 μl reactions containing 0.2 μM primers, 1× buffer, 0.2 mM dNTPs, 2.5 U Platinum *Taq* DNA polymerase and 3 mM MgCl_2_. Primer sequences are detailed in Table 1. PCR cycling conditions were as follows: 95 °C for 3 min followed by 35 cycles of 30 sec each at 95 °C; 58 °C, 72 °C for 30 sec, and a final elongation step of 5 min at 72 °C. PCR products were purified using the Wizard clean up kit (Promega) and cloned into the pJET1.2/blunt vector (Fermentas). Insertion of PCR products was verified by digestion with *Bgl*II and positive clones were sequenced using conventional Sanger sequencing (Eurofins Genomics). Combined bisulfite restriction analysis (COBRA) was carried out using *Taqα*I, and/or *Aci*I as outlined in ^27^. Sequence analysis and alignment was performed using DNAStar EditSeq, MegAlign (www.dnastar.com) and BiQ Meth Analyzer (http://biq-analyzer.bioinf.mpi-inf.mpg.de). During this analysis, sequences with low C-T conversion rate (< 95%) and with a high number of sequencing errors (sequence identity with genomic sequence less than 80%) were excluded from the alignment. Identical clones were also excluded from the analysis.

### Illumina sequencing and initial quality control

Pooled libraries were sequenced at the Michigan State University Research Technology Support Facility. Paired-end reads (2 × 150 bp) were obtained by Illumina sequencing of each pooled library on four lanes of a HiSeq 2500 sequencer, in rapid run mode. After pooling data from all lanes, bisulfite-treated (BS) libraries yielded 45.3–67.4 million read pairs per sample; comparably, non-bisulfite (NON-BS) libraries yielded a total of 40–50 million read pairs per sample across all lanes (Supplementary Table 1).

Quality control of raw read pairs using *FastQC* (www.bioinformatics.babraham.ac.uk/projects/fastqc) revealed similar QC metrics for both infected and control samples (Supplementary Table 2). Although samples from animals 1, 4, 5, and 6 raised warnings of *Over-Represented Sequences*, this warning was systematically triggered by N-polymers in the second mate, a technical issue resolved by quality trimming. As expected, BS libraries raised significantly more *Per Base Sequence Content* and *Per Base GC Content* than NON-BS libraries, due to the nature of the bisulfite treatment (Supplementary Table 2).

### Adapter and quality trimming

Stringent adapter trimming (overlap ≥1 bp at the 3’ end of each read), and quality trimming (Phred ≤ 20 from the 3’ end of each read) using *Trim Galore!* [version 0.4.1] (www.bioinformatics.babraham.ac.uk/projects/trim_galore) left 97.5–98.4% of raw read pairs in bisulfite-treated samples, and 93.1–94.3% of raw read pairs for NON-BS samples. However, this trend was reversed at the nucleotide level, with 77.5–86.9% of sequenced bases left in BS libraries, against 84.8–87.3% in NON-BS libraries. Notably, BS libraries generally showed higher levels of adapter contamination (54.2–77.3% of raw reads) relative to NON-BS libraries (48.6–58.2% of raw reads), based on the stringent detection rule described above. Second read mates displayed a larger proportion of low-quality sequenced bases trimmed (7.6–13.5% of raw sequenced bases) relative to first mates (1.9–4.5% of raw sequenced bases), in both BS and NON-BS libraries (Supplementary Table 1).

Notably, quality control of trimmed libraries revealed a significant improvement of *Over-Represented Sequences*, and full resolution of *Adapter Content* warnings (data not shown). As a result of stringent adapter trimming (even a single trailing A at the 3’ end was trimmed; following Trim Galore! default settings), all samples raised warnings of *Per Base Sequence Content* caused by the severe under-representation of A nucleotides at the 3’ end of reads, a known artefact of the stringent trimming process with no notable repercussion on the subsequent alignments and methylation calls.

### Alignment of bisulfite-treated libraries

BS libraries were aligned using *Bismark* [version 0.15.2] ^28^ and the *Bowtie2* aligner [version 2.2.6] ^29^ in strand-specific (directional) mode to computationally generate bisulfite-converted copies of the top and bottom strands of the *Bos taurus* UMD3.1 genome assembly ^30^. Alignment efficiency (*i.e.*, read pairs aligned to a unique locus) reached 59.7–68.3%, for a total of 28.8–42.7 million read pairs aligned uniquely per sample. Aligned reads were found evenly distributed between the top and bottom strands of the BS-converted genome (Supplementary Table 3). Bismark methylation calls revealed methylation levels in the range of 69.2–73.8% in CpG context, for a total of 113–156 million methylation calls per sample. In contrast, non-CpG context displayed markedly low methylation levels (0.7–1.7%), with orders of magnitude larger counts of methylation calls, owing to their broader definition of methylation context (392–543 million calls in CHG context; 1.1–1.5 billion calls in CHH context: H corresponds to A, T or C).

### Deduplication of aligned bisulfite-treated libraries

Paired-end alignments where both mates aligned to the same position in the genome were removed from the Bismark alignment output using the *deduplicate_bismark* script to mitigate the impact of duplicate DNA fragments sequenced. This procedure discarded 6.1–18.8% aligned read pairs, leaving 25.5–38.7 million aligned read pairs for subsequent methylation calls (Supplementary Table 4).

### Methylation calls

The *bismark_methylation_extractor* script was used in a two-pronged approach. First, methylation calls extracted from the full sequence of aligned read pairs were used to evaluate *M*-bias across the aligned mates. *M*-bias plots show the methylation proportion across each possible position in the read, and reveal anomalies at any position of the sequenced reads, often found toward ends of the sequenced reads. After analysis of the *M*-bias plots generated in the first pass, the second call to the *bismark_methylation_extractor* script was set to ignore the first seven bases at the 5’ end of both read mates. Collated reports of the Bismark pipeline leading to the final methylation calls are available as HTML files in Supplementary File 1.

### Statistical analyses

Methylation calls in CpG context were imported from the individual Bismark CpG reports, combined and processed in a *BSseq* container of the *bsseq* Bioconductor package [version 1.9.2; www.bioconductor.org/packages/bsseq] ^31^. Genomic coordinates of CpG islands on all sequences (*i.e.*, unmasked sequence CpG island track) in the UMD3.1/bosTau6 assembly were obtained from the UCSC Table Browser ^32^.

### Non CpG methylation

Non-CpG methylation (CHH and CHG) present in the WGBS reads were analysed using the methylKit R package ^33^. De-duplicated bam files produced during alignment with Bismark were sorted and saved as sam formatted files. Individual CHH or CHG were imported separately. These files were imported to the methylKit object using strict criteria: at least 10× coverage per feature and the feature must be present across all samples. This resulted in a total of 685,311 and 284,641 features for CHH and CHG respectively, with a median coverage of 40× for CHH and 41× for CHG. Median coverage between all samples was used to calculate a scaling factor to normalize the coverage across samples. Differential methylation was determined for individual features with an overdispersion parameter included (shrinkMN) and Benjamini-Hochberg (B-H) FDR adjustment for multiple comparisons ^34^. All downstream analyses were carried out separately for CHH and CHG methylation. Average methylation between groups was tested using a paired *t*-test.

### Expression dynamics of genes associated with chromatin configuration and DNA methylation

For the bAM samples used for the WGBS analysis at 24 hpi described here, differentially expressed genes were previously identified in the same *M. bovis*-infected bAM, relative to the non-infected bAM at 2, 6, 24 and 48 hpi using RNA-seq ^23^. A comprehensive list (EPI-list) of 151 genes previously identified as being involved with histone modifications or DNA methylation was generated from the literature (Supplementary File 2). RNA-seq transcriptomics data was mined, at each time point, using the EPI-list. Ultimately, 86 genes were identified from the RNA-seq data (Supplementary File 2). These 86 genes were denoted as genes of interest (GOI) and their expression was determined at 2, 6, 24 and 48 hpi using the previously published lists of differentially expressed genes (*P* < 0.05, B-H FDR-adjusted).

### Gene ontology enrichment analyses

Gene ontology enrichment analyses were performed using the Bioconductor *GOseq* software package ^35^ and the annotation package *org.Bt.eg.db* (https://bioconductor.org/packages/org.Bt.eg.db). Notably, the probability weighting function (PWF) supplied to *GOseq* was calculated without length bias for the analysis of promoters, as those were defined in this study to a constant width of 2 kb (1.5 kb upstream and 500 bp downstream of TSS).

## Results

### WGBS summary statistics

In summary, 16 individually barcoded WGBS libraries, prepared using bAM DNA extracted from eight *M. bovis*-infected and eight non-infected samples, were sequenced on an Illumina HiSeq 2500 sequencer in rapid run mode. This generated 45.3–67.4 million read pairs per sample and ∼32× sequencing depth per condition (*M. bovis*-infected and non-infected bAM). These data satisfy previously defined criteria for WGBS, with respect to the number of independent biological replicates and the sequencing depth (www.roadmapepigenomics.org/protocols) ^36^. An assessment of bisulfite conversion rates was performed using non-CpG methylation according to Clark *et al.* ^37^. Based on this approach, conversion efficiencies were > 99% using CHH methylation values and > 98% using CHG methylation values.

When all CpG dimers in the reference bovine genome (UMD3.1) were considered— approximately 55.1 million stranded loci, including potentially unmappable CpG dimers—the 25.5–38.7 million aligned read pairs used for methylation calls led to an average 1.1–1.4 methylation call per individual strand-specific CpG dimer in each individual sample. As a result of collapsing methylation calls as unstranded CpG loci, the even distribution of aligned read pairs on both strands of the reference genome (Supplementary Table 3) doubled coverage to 2.12–2.78× per unstranded CpG dimer (∼27.5 million unstranded loci). Unstranded methylation calls were used from this point onwards. While the mean coverage of CpG dimers covered in at least one sample was similar (2.13–2.81×; ∼27.3 million loci), CpG dimers covered in all samples was larger for each sample (4.6–5.9×; 4.7 million loci). Notably, the average coverage of all CpG dimers in known CpG islands (CGIs)—including CpG dimers with null coverage— was similar to those latter values (3.9–5.4×; 2.9 million loci), suggesting a consistent coverage of CGIs across all samples.

### Genome-wide scan for differentially methylated regions 24 hpi

An unbiased genome-wide scan was performed to identify potential differentially methylated regions (DMRs), including only CpG loci with at least two methylation calls for at least six of the eight biological replicates in each sample group, thereby ensuring at least 12× coverage for any CpG dimer in both sample groups. As a comparison, the analysis was repeated after randomising samples from both infection groups to produce a distribution of *t*-statistics under the null hypothesis. The *bsseq* package was used to calculate *t*-statistics in a paired design for both original and randomised sets of sample (Fig. 2A and 2B).

**Figure 2.**
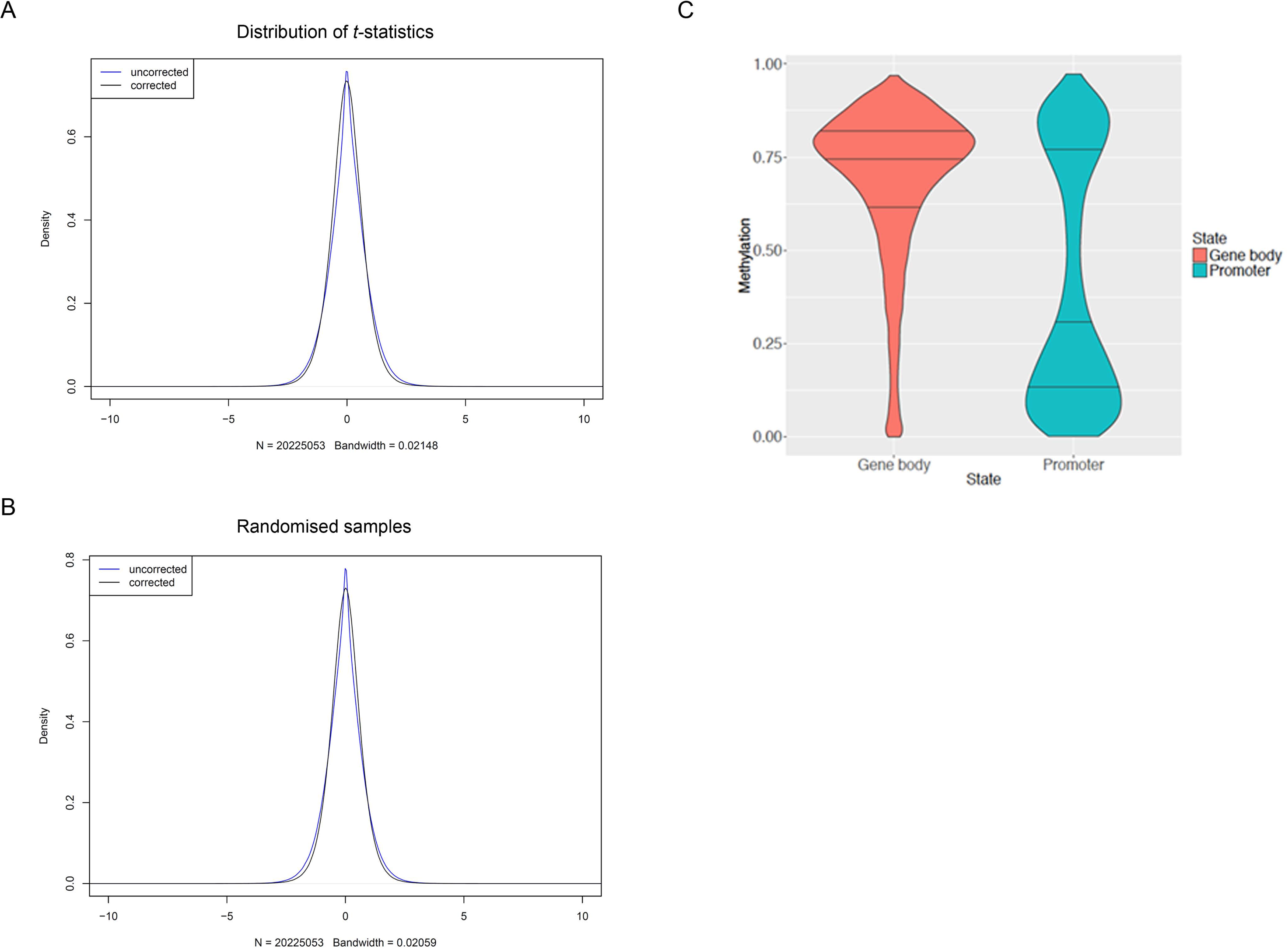
**(A)** Distribution of paired *t*-statistics between *M. bovis*-infected and control non-infected bAM samples based on smoothed WGBS data. **(B)** Distribution of paired *t*-statistics between randomised samples based on smoothed WGBS data. **(C)** Distribution of average methylation level (%) in promoters and gene bodies across all samples. Only CpG loci with coverage greater or equal to 10 were considered. Only genes where gene body and promoter both had 10 or more sufficiently covered CpG loci were considered.

Potential DMRs were identified as genomic regions including at least three loci with absolute *t*-statistics greater than 4.6 and a mean difference in methylation level (across samples and loci) greater than 10% between the two groups. This analysis did not reveal significant differences in methylation between infected and non-infected bAM; regions identified in the original data were comparable to randomised data in number of regions identified and their properties (*e.g.*, width, number of methylation loci, sum of *t*-statistics, proportion of regions showing increased and decreased methylation level) (Supplementary Table 5).

### Distribution of methylation across different genomic regions

Following the genome-wide scan, DNA methylation was determined at the following defined functional genomic elements: gene bodies, intergenic sequences and proximal promoters. Genomic elements were defined as described previously by Peat and colleagues ^38^ and the number of regions compared in each category are outlined in Table 2. As expected for a differentiated/somatic cell type, the majority of CpGs within intragenic sequences, gene bodies and CpG-deficient promoters were widely methylated ^39,40^. CpG-rich promoters containing a CpG island (CGI) or overlapping a CGI (Promoter CGI and CGI promoter) were mostly hypomethylated (Fig. 3). Interestingly, CGIs remote from annotated gene promoters (non-promoter CGIs) showed variable methylation—most were hypermethylated (>75% methylated, 18,581 CGIs) with 9,103 non-promoter CGIs hypomethylated (<25%).

**Table 2.** Number of tiles/regions in each category.

| **Region** | **Control** | ***M. bovis*** |
| --- | --- | --- |
| **Intergenic** | 347,561 | 347,561 |
| **Gene Body** | 166,466 | 166,466 |
| **CGI Promoters** | 12,047 | 12,047 |
| **Non-CGI Promoters** | 13,413 | 13,413 |
| **Promoter CGIs** | 11,222 | 11,222 |
| **Non-Promoter CGIs** | 30,284 | 30,284 |

**Figure 3.**
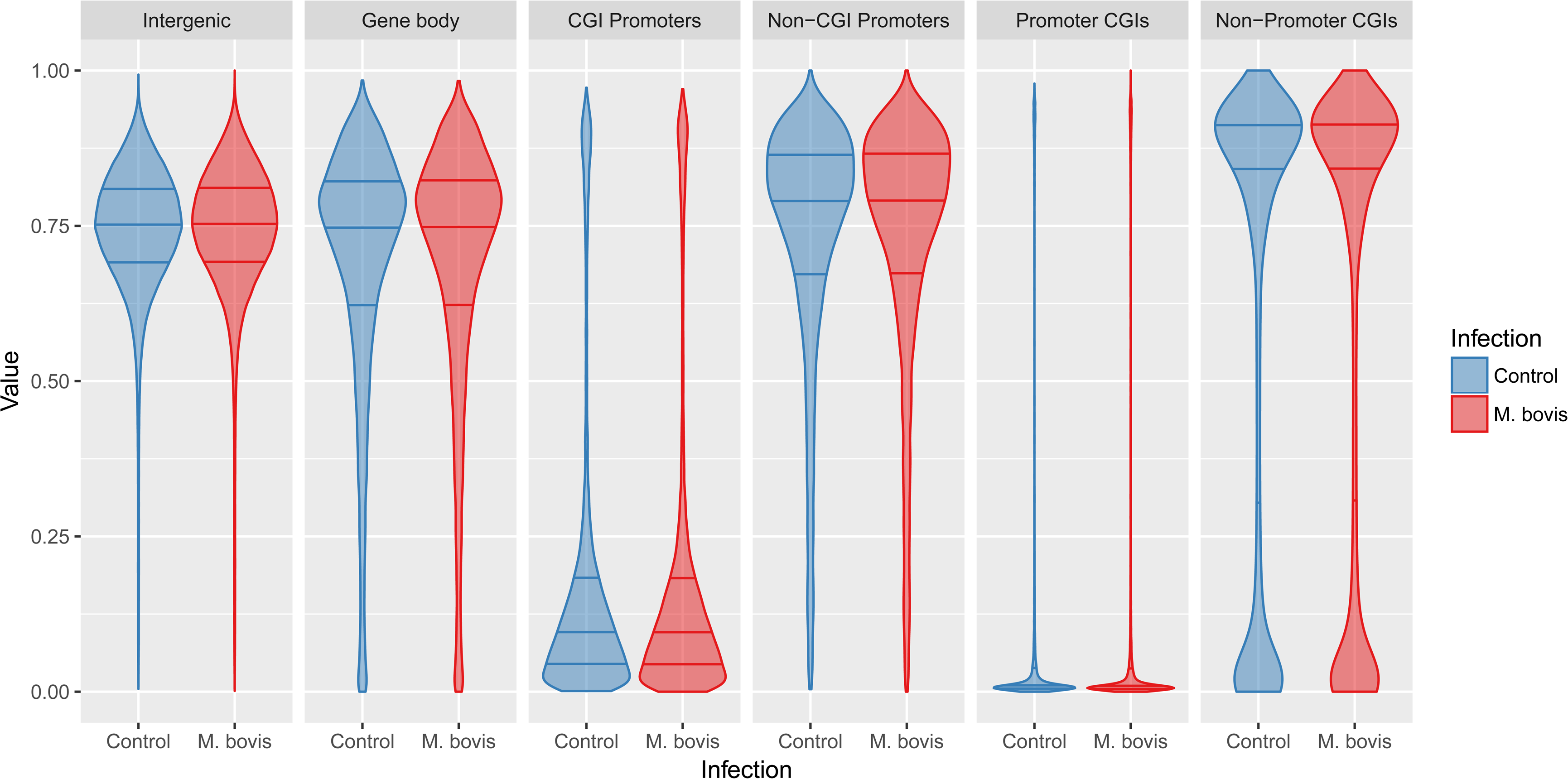
Distribution of DNA methylation in different genomic contexts in non-infected and *M. bovis*-infected bovine alveolar macrophages (24 hpi) Analysis of WGBS data from *M. bovis*-infected and non-infected bovine alveolar macrophages (bAM) revealed that genomic methylation, in the context of CpGs, was not altered at any of the sequence features outlined (intergenic regions, gene bodies, or promoters with or without CpG islands (CGIs) in the host following infection. Blue and red violins represent non-infected and *M. bovis*-infected bAM, respectively.

### Pyrosequencing validation of WGBS at key immune function genes

To confirm the WGBS observation that DNA methylation was not different between control and infected bAM, 24 hpi, a small panel of key immune genes, *TNF, IL12A, TLR2, NFKB2, C1QB, NOS2* and *DTX4* were selected for targeted analysis by pyrosequencing. Transcription of these genes has previously been shown to be upregulated in bAM 24 hpi with *M. bovis* ^21,23^; the specific loci that were analysed by pyrosequencing are detailed in Fig. 4. Four of the loci are hypomethylated (*TNF, IL12A, TLR2* and *NFKB2*), one is intermediately methylated (*NOS2*) and two are highly methylated (*C1QB*). Using DNA isolated from a parallel set of control (*n* = 4) and infected (*n* = 4) bAM samples, average methylation levels at the proximal promoter regions of *NFKB2, TLR2, IL12A* and *TNF*, and the gene bodies of *C1QB, NOS2* and *DTX4*, were determined. Statistical analysis using a paired *t*-test did not reveal significant differences (*P* ≥ 0.05) in mean methylation levels between the examined loci of infected and non-infected bAM samples (Fig. 5), supporting the WGBS observation that *M. bovis* does not have an effect on the CpG methylation in bAM.

**Figure 4.**
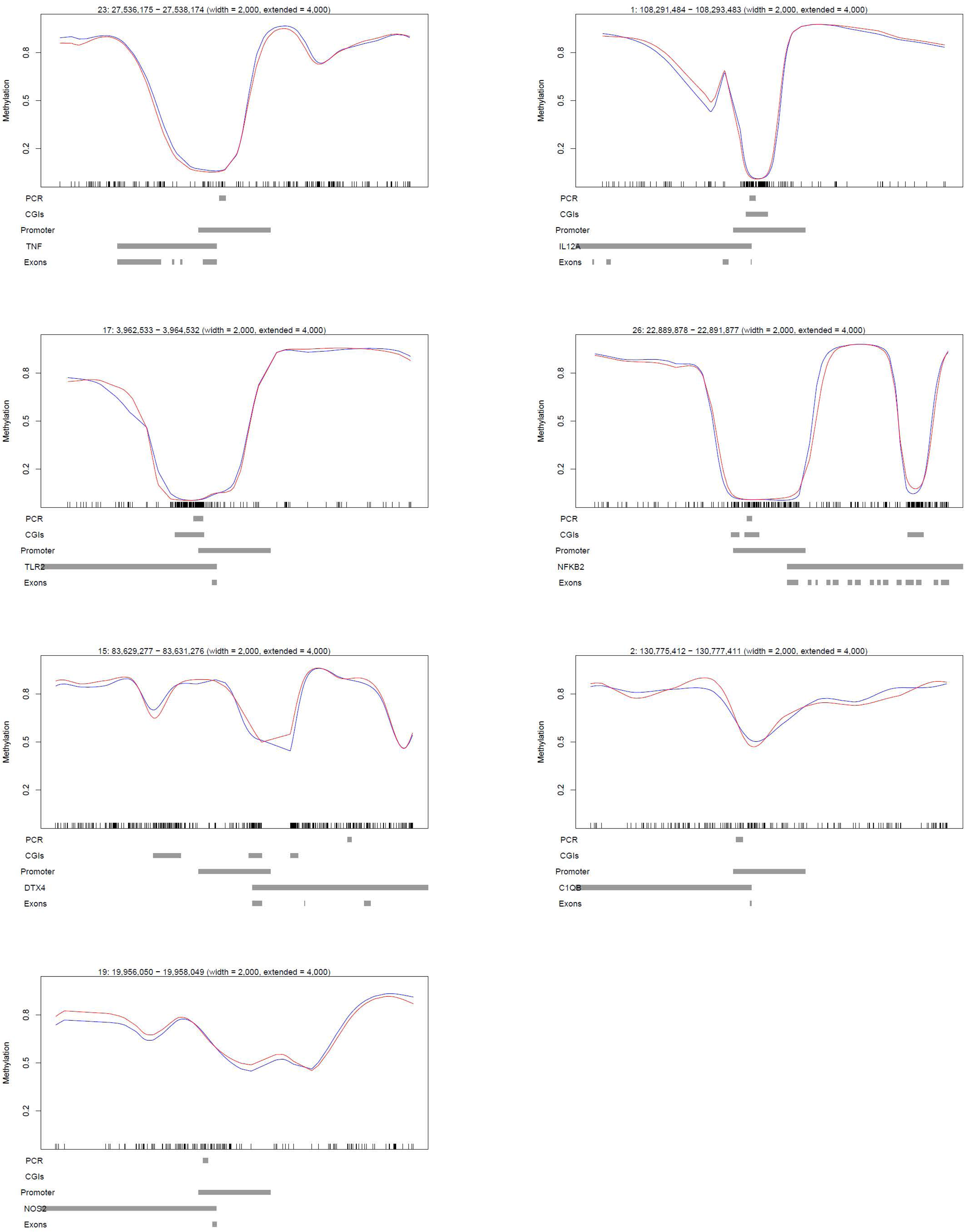
Schematic representation of WGBS data at loci related to immune function. WGBS proximal promoter plots. Representative plots showing the average methylation spanning a 10 kb region at the 5’ end of the *TNF, IL12A, TLR2, NFKB2, DTX4, C1QB* and *NOS2* genes. The red and blue lines represent average methylation levels for infected and control samples, respectively.

**Figure 5.**
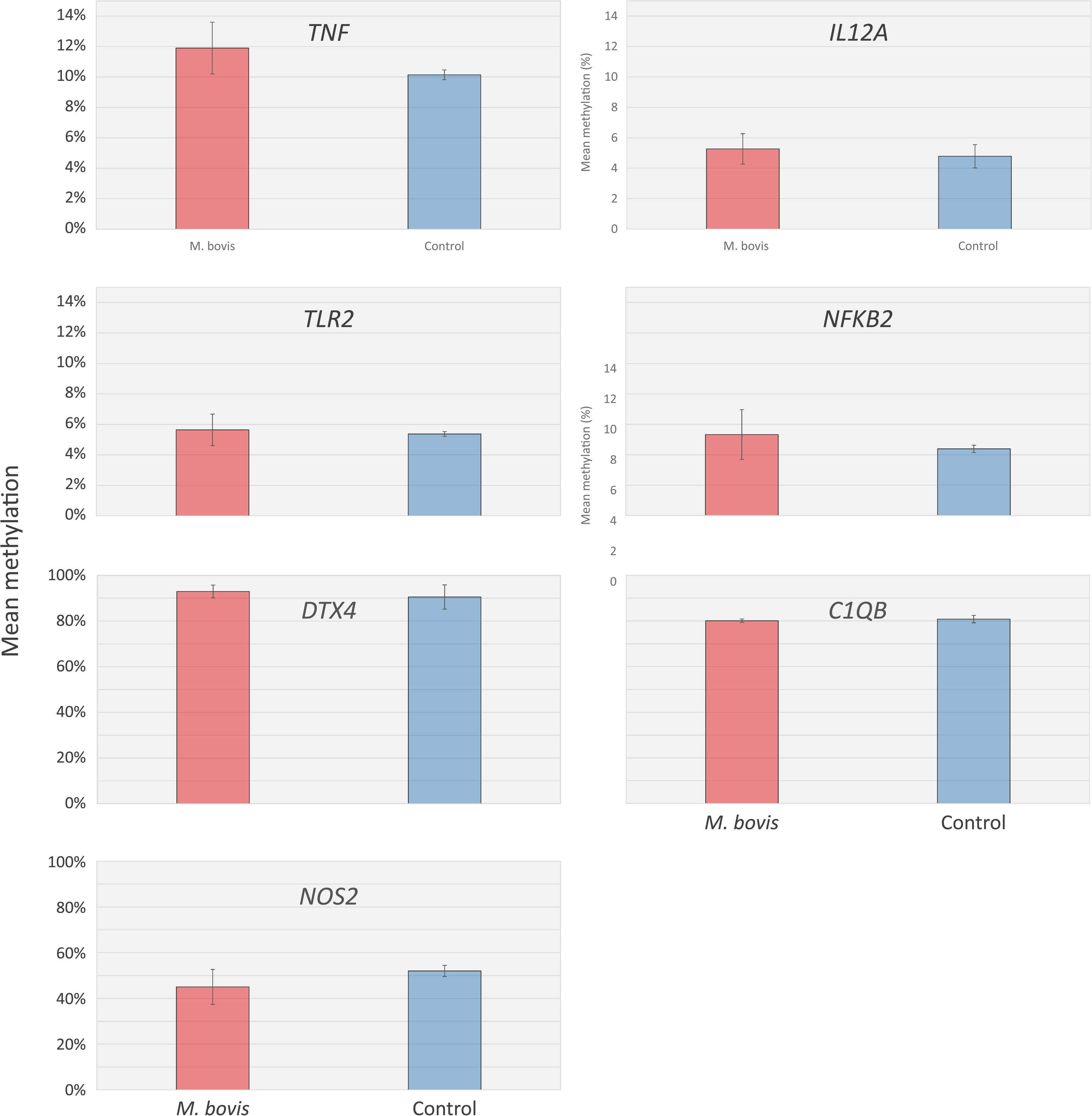
Pyrosequencing validation of whole-genome bisulfite sequencing (WGBS) results. Locations of the pyrosequencing assays are denoted by ‘PCR’ in Fig. 4. Methylation was not different at any of the loci tested (paired *t*-test *P* ≥ 0.05) between *M. bovis*-infected and non-infected control bAM. The number of CpG dinucleotides analysed at each loci were *TNF* (9 CpGs), *IL12A* (11 CpGs), *TLR2* (9 CpGs), *NFKB2* (8 CpGs), *C1QB* (4 CpGs), *DTX4* (3 CpGs) and *NOS2* (1 CpG).

### Promoter methylation level and gene ontology in bovine alveolar macrophages

Leveraging the absence of significant DMRs between infected and non-infected bAM samples, methylation calls were pooled across all sixteen samples (eight *M. bovis*-infected and eight controls) to analyse methylation levels in bAM gene promoters, with maximal coverage. In this analysis, promoters were defined as regions spanning 1.5 kb upstream and 500 bp downstream of each transcription start site (TSS), with a minimum of 10 CpGs each associated with at least five methylation calls were included (26.8 million loci). Of the 24,616 genes annotated in the bovine genome (Ensembl BioMart March 2016 archive), 22,964 were retained for this analysis, on the basis that their promoter contained at least ten loci, each locus having at least five methylation calls. For those genes, mean promoter methylation was estimated and summarised, alongside average gene body methylation, in Fig. 2C.

Notably, the vast majority of gene promoters were found at either extreme of the methylation range. Indeed, 18,438 promoters (80.3%) display methylation levels greater than 75% or lower than 25% (8,145 ≥ 75% methylated; 10,293 ≤ 25% methylated). However, 2,580 promoters (9.7%) displayed an average intermediate methylation level (IM, 33–66%). Strikingly, gene ontology (GO) analysis of the genes associated with IM promoters (33–66%) using the *GOseq* package ^35^ revealed a marked enrichment for immune-related GO categories including “defense response” (*P* < 10^−08^), “defense response to bacterium” (*P* < 10^−07^), “response to bacterium” (*P* < 10^−07^), “chemokine-mediated signaling pathway” (*P* < 10^−06^) and “chemokine activity” (*P* < 10^−04^), among others (Supplementary Table 6). In contrast, no significant enrichment for immune-related GO categories was found for promoters with methylation levels 0–1% (759 promoters), 0–10% (5,605), 10–20% (3,255), or 90–99% (1,997) (Supplementary Table 6). Instead, the latter only suggested enrichment for generic GO categories (*e.g.*, “intracellular organelle”, “transcription regulatory region DNA binding”).

A hallmark of some imprinted genes is that they contain a 5’ differentially methylated region that is IM (resulting from parent-of-origin specific methylation patterns); therefore, the IM promoter list was interrogated for known bovine imprinted genes (www.geneimprint.com/site/genes-by-species.Bos+taurus). This analysis confirmed IM at the promoters of the following imprinted genes; *PLAGL1, SNRPN, MEST, PEG10, GNAS* and *NNAT* (Supplementary Fig. 1).

### Targeted analysis of intermediately methylated (IM) gene promoters

To confirm the presence of IM at immune gene promoters, COBRA and clonal analysis of bisulfite PCR products was performed. Firstly, proximal promoter alignment plots for the IM group were visually screened to remove promoters that were included due to averaging of sequences with high and low methylation (example of this in Fig. 6). This analysis was restricted to IM promoters containing a minimum of 30 CpGs (1,034 loci), to ensure sufficient CpG coverage during COBRA and clonal bisulfite sequencing analysis. Of the 1,034 IM promoters, 267 promoters remained in the IM group and 60/267 (22.5%) of them had a promoter CGI (Supplementary File 3). GO analysis of these 267 IM gene promoters with ≥ 30 CpGs revealed enrichment for NADH dehydrogenase-associated activity (Supplementary Table 6). Two immune-related gene promoters with the highest CpG content, *C1QB* and *IL2RA*, were selected for further analysis (Fig. 7). Clonal analysis revealed that, although there are clearly hypermethylated and hypomethylated *C1QB* and *IL2RA* alleles, the prominent allelic methylation pattern is mosaic (Figs. 8 and 9); suggesting that the IM promoters analysed are methylated in an allele-independent as opposed to an allele-specific pattern, an observation that has been previously reported ^41^. To further confirm our WGBS and clonal bisulfite sequencing results we carried out COBRA on *C1QB* and *IL2RA* IM regions, using *M. bovis*-infected and non-infected bAM (Fig. 8 and 9). Results from COBRA support our observation that the *C1QB* and *IL2RA* proximal promoters were IM. Additionally, bovine sperm, kidney, liver and heart samples were assessed using COBRA to determine whether IM might be tissue-specific. COBRA indicated that *IL2RA* was almost completely methylated in sperm and predominantly methylated in the kidney, liver and heart; suggesting a potential tissue-specific IM in bAM (Fig. 9). This possible tissue-specific IM pattern was not observed at the *C1QB* locus (Fig. 8).

**Figure 6.**
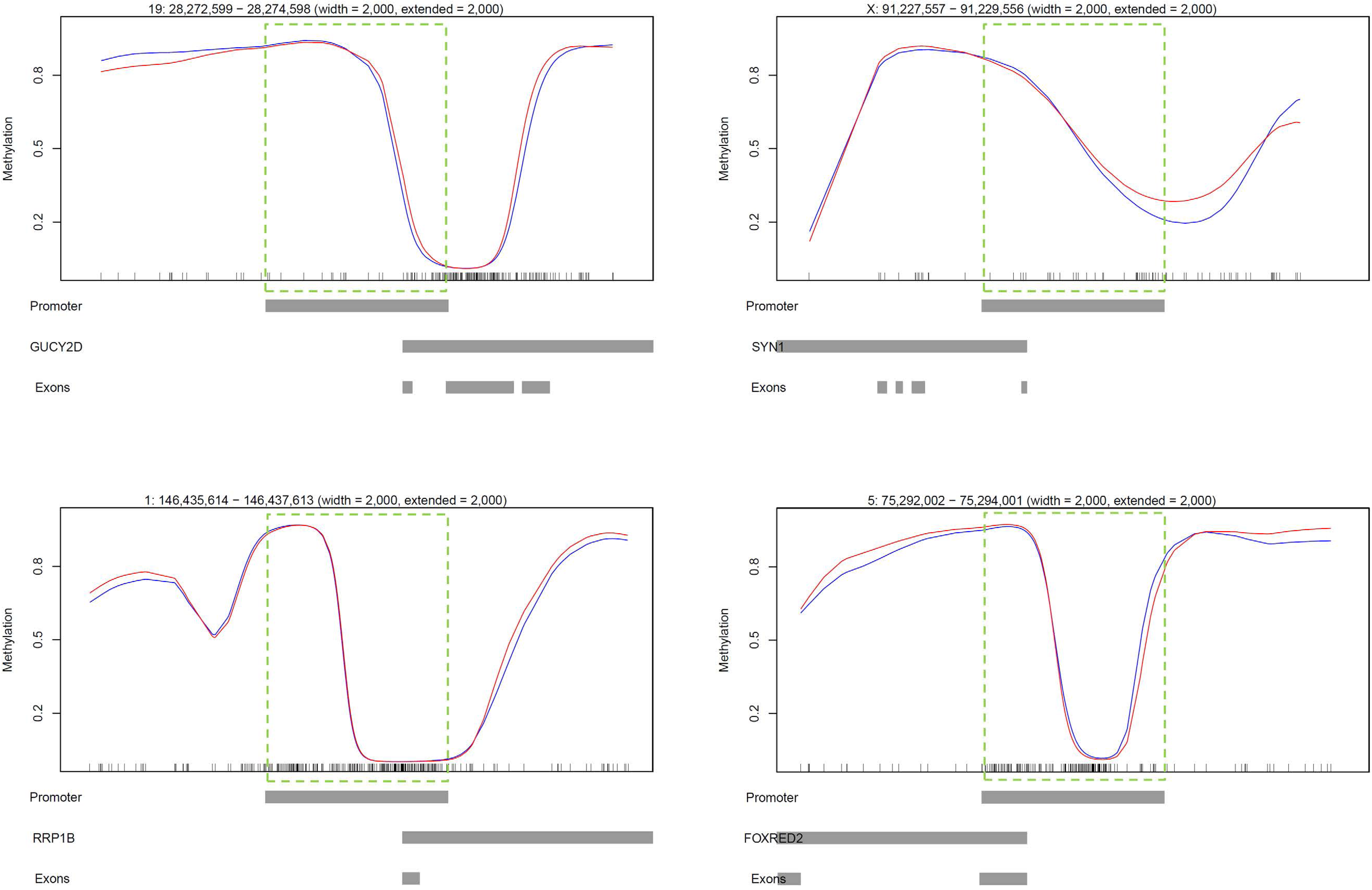
Analysis of promoters with highly methylated and unmethylated sequence. 1,034 proximal promoters, with a minimum of 30 CpGs, shown to be intermediately methylated (IM) in the WGBS analysis were visually inspected to remove false positives. 767 were eliminated from the IM group due to averaging of highly methylated and unmethylated CpGs in the proximal promoter region (green dashed box). Four examples are presented here: the *GUCY2D, SYN1, RRP1B* and *FOXRED2* genes. Red and blue lines represent average methylation levels for infected and control samples, respectively.

**Figure 7.**
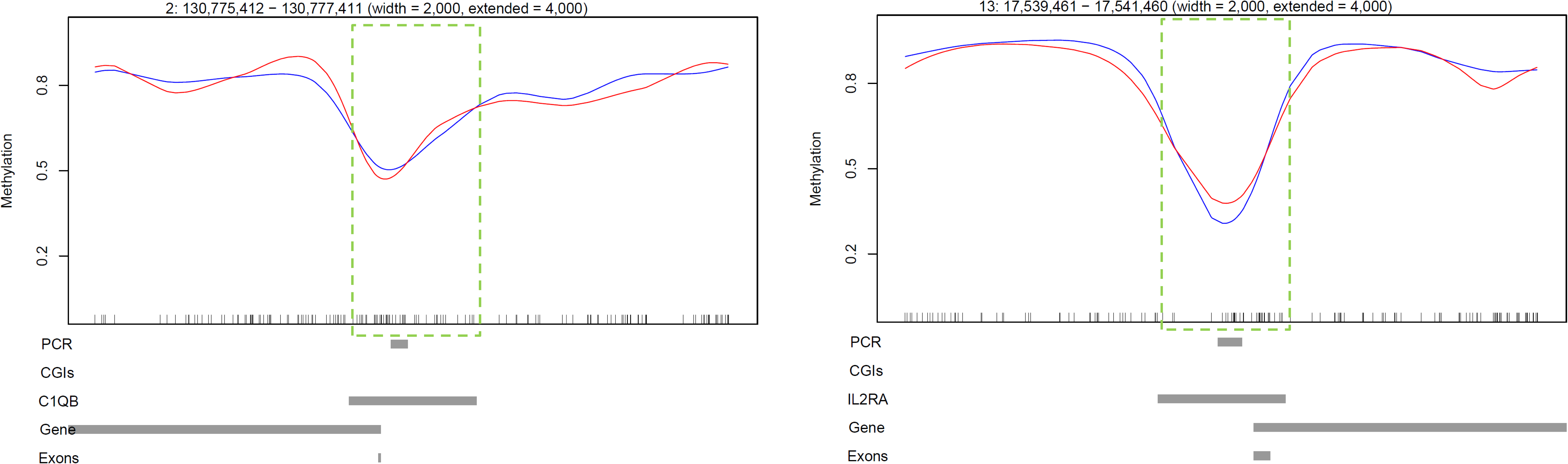
Gene promoters for combined bisulfite restriction analysis (COBRA) and clonal analysis. WGBS aligments at the *C1QB* and *IL2RA* gene loci. Each panel represents a 10 kb region at the 5’ end of the gene. Green dashed boxes illustrate the *C1QB* and *IL2RA* proximal promoter regions (the TSS minus 1.5 kb, plus 500 bp) identified as intermediately methylated (IM) during WGBS data analysis (average methylation 33–66%). PCR: region analysed using bisulfite PCR, cloning and Sanger sequencing; CGIs: CpG islands; Gene: transcribed region; Exons: shows the location of the first exon; Red line: *M. bovis*-infected bAM; Blue line: non-infected control bAM.

**Figure 8.**
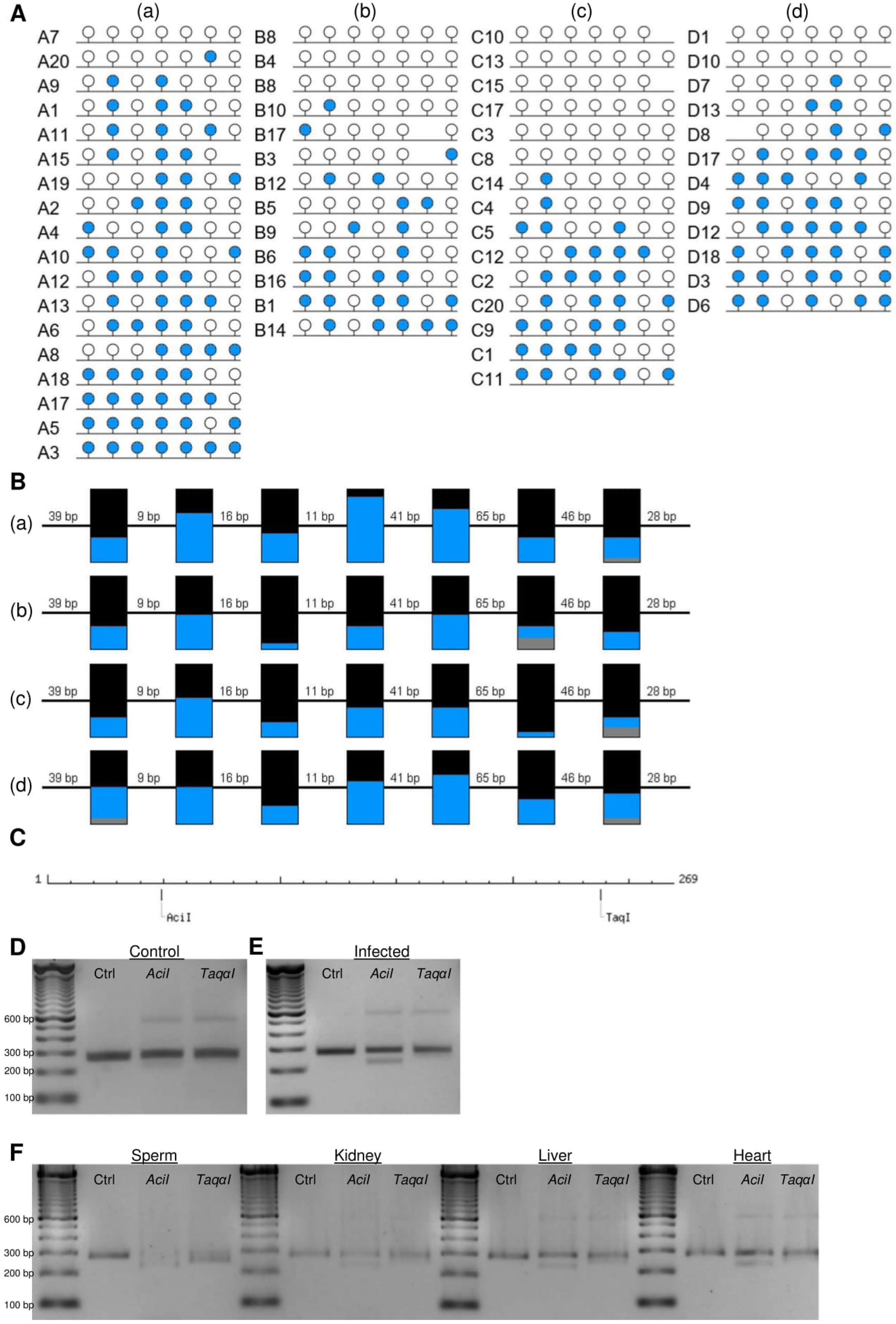
Confirmation of an intermediately methylated promoter region at the *C1QB* gene locus. **(A)** Clonal analysis of seven CpG dinucleotides in a 269 bp fragment of the bovine *C1QB* 5’ promoter region, *a*–*d* represent sequencing of four biological replicates. Closed and open circles denote methylated and unmethylated CpGs, respectively. **(B)** Aggregated representation of methylation status at CpGs 1–7 in the *C1QB* proximal promoter region; (a)-(d) represent animals A-D, numbers between boxes indicate genomic distance between CpGs while numbers above boxes indicate the position of the CpG within the analysed region; BLUE = methylated, BLACK = unmethylated, GREY = not present; **(C)** Schematic representation of the analysed C1QB region and the recognition sites of *Aci*I and *Taqα*I as obtained by NEBcutter V2.0; length is displayed in bp; **(D) – (F)** COBRA results of Uninfected **(D)**, Infected **(E)** and tissue samples **(F)** digested with AciI, TaqαI or undigested (Ctrl).

**Figure 9.**
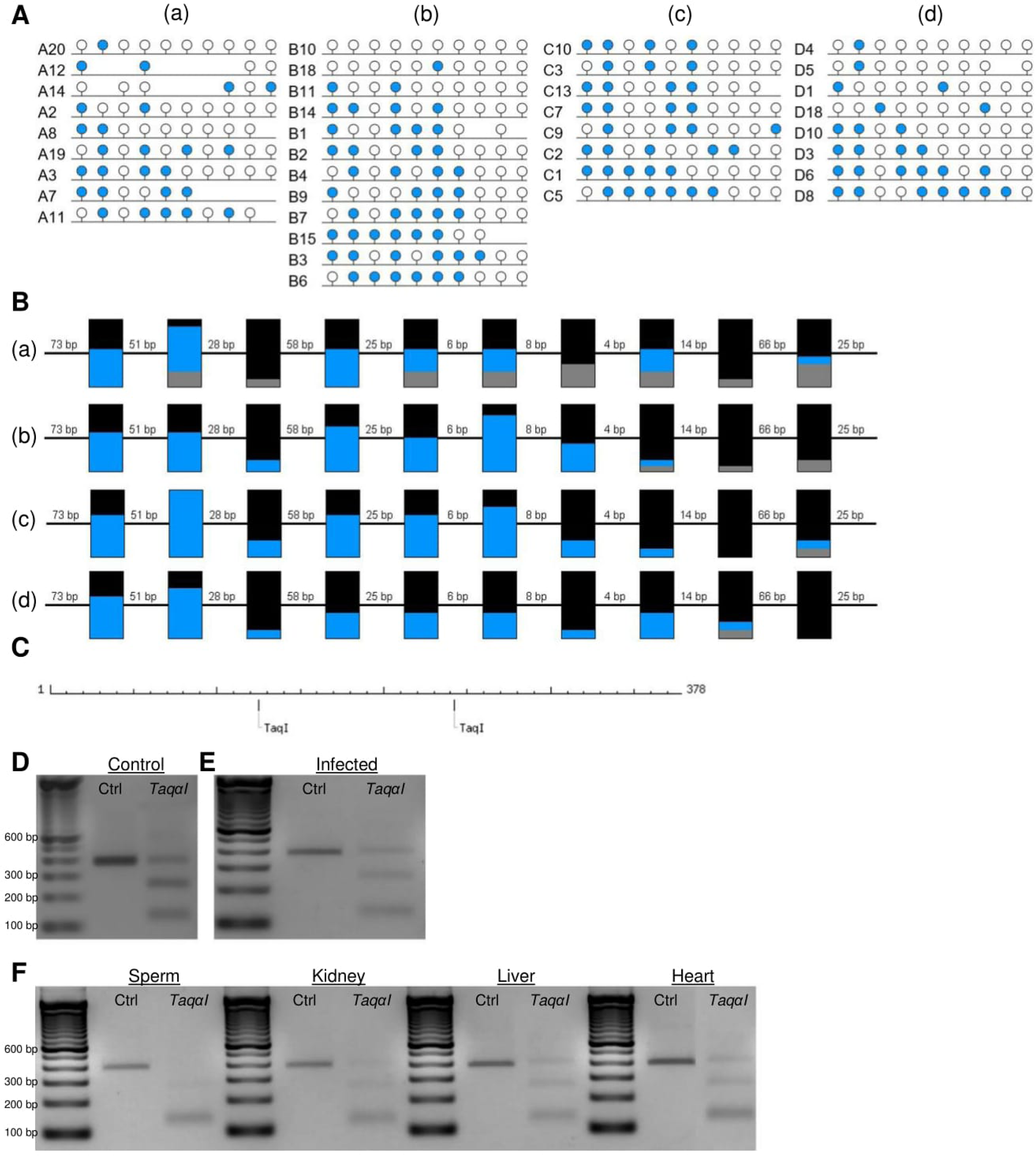
Confirmation of an intermediately methylated promoter region at the *IL2RA* gene locus. **(A)** Clonal analysis of 10 CpG dinucleotides in a 378 bp fragment of the bovine *IL2RA* 5’ promoter region, *a*–*d* represent sequencing of four biological replicates. Closed and open circles denote methylated and unmethylated CpGs, respectively. **(B)** Aggregated representation of methylation status at CpGs 1–10 in the *IL2RA* proximal promoter region; *a*-*d* represent animals A-D, numbers between boxes indicate genomic distance between CpGs while numbers above boxes indicate the position of the CpG within the analysed region; BLUE = methylated, BLACK = unmethylated, GREY = not present; **(C)** Schematic representation of the analysed *IL2RA* region and the recognition sites of *Taqα*I as obtained by NEBcutter V2.0; length is displayed in bp; **(D) – (F)** COBRA results of Uninfected **(D)**, Infected **(E)** and tissue samples digested with *Taqα*I or undigested (Ctrl).

### Non-CpG methylation analysis

Overall, we found a low level of methylation in the context of CHH: mean values of 0.98% and 0.96% were estimated for control and *M. bovis*-infected bAM, respectively (Fig. 10). CHH methylation was not different between control and infected-bAM (*t*-statistic 1.32, df = 12.7, *P* > 0.05). Similarly, for CHG methylation we found an overall low mean methylation of 1.53% and 1.49% for control and *M. bovis*-infected bAM, respectively (Fig. 10). There was no difference for this mean methylation in CHH context between groups (*t*-statistic 1.26, df = 13.5, *P* > 0.05). Neither clustering on all data nor top 5,000 most variable features revealed any patterns in these data sets for CHH and CHG methylation. This was also concordant with differential methylation tests showing no loci as significantly differentially methylated between groups by a methylation difference greater than 1% and q-value = 0.01.

**Figure 10.**
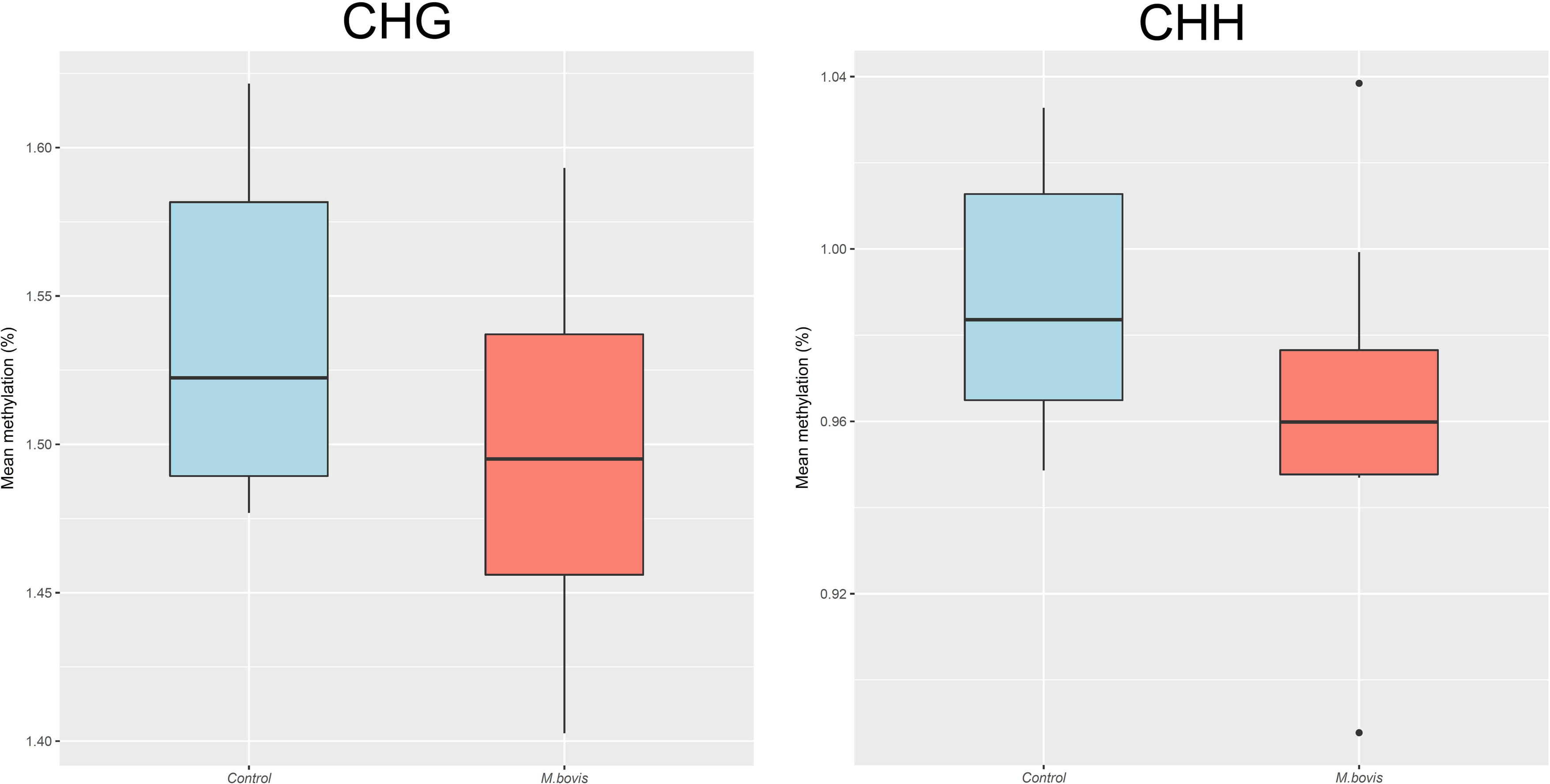
Non-CpG methylation levels differ but not significantly in bovine alveolar macrophages (bAM) infected with *Mycobacterium bovis*. Mean methylation at the non-CpG contexts CHG (left) and CHH (right) for control and *M. bovis*-infected bAMs, respectively: differences were not significant by *t*-statistic.

### Relationship between *M. bovis* infection and expression of chromatin and DNA modifiers

Based on our WGBS results, bAM DNA methylation is not affected by infection with *M. bovis* at 24 hpi; therefore, we next determined whether *M. bovis* infection has an effect on chromatin. To do this, transcription analysis of chromatin and DNA modifying enzymes was carried out using our previously published RNA-seq data from *M. bovis*-infected bAM ^23^ and a similar approach to that detailed by Nestorov and colleagues ^42^. A list of 151 genes (EPI-list) that encode chromatin and DNA modifying enzymes was assembled from the literature (Supplementary File 2). To identify chromatin and DNA modifying-associated genes that were detected by RNA-seq, differentially expressed genes (*P* < 0.05, B-H FDR-adjusted) at each time point were compiled and searched using the list of 151 known genes. This identified a list of 86 genes of interest (GOI). The number of GOIs was determined at each time point and the results were as follows: 2 hpi 0/86, 6 hpi 8/86 (3 upregulated, 5 downregulated), 24 hpi 37/86 (16 upregulated, 21 downregulated) and 48 hpi 48/86 (19 upregulated, 29 downregulated) (Supplementary File 2). *HDAC5, KDM2B, EZH1, PRDM2, SETMAR, SMYD4* and *USP12* were differentially expressed at all time points post-infection (excluding 2 hpi).

## Discussion

Here we present genome-wide DNA methylation profiles of bAM infected with *M. bovis* versus non-infected controls at 24 hpi. We show that CpG methylation in bAM is not altered in response to *M. bovis* at 24 hpi. Since previous studies suggest that DNA methylation changes are established relatively late in the silencing pathway and are preceded by alterations to histone modifications and chromatin packing ^43^, our results may reflect the early post-infection time point examined in this study. Examination of the WGBS data, focusing on the relationship between DNA methylation and proximal promoters, revealed an enrichment of gene promoters that were intermediately methylated.

Global methylation patterns were first analysed using an unbiased genome-wide scan to identify differentially methylated loci between *M. bovis*-infected and control bAM. Following this, we examined the impact of infection on the bAM methylome in greater detail by assessing DNA methylation at specific genomic features. Given the relationship between promoter methylation, gene body methylation and transcription ^44^, these genomic features comprised the main focus of these analyses. Promoters and CGIs were separated into the following categories as previously described ^38^: CGI promoters (*i.e.*, gene promoters overlapping a CpG island), non-CGI promoters, promoter CGIs and non-promoter CGIs. None of these approaches revealed any differentially methylated loci between *M. bovis*-infected and non-infected control bAM at 24 hpi. Therefore, it is unlikely that the substantial transcriptomic perturbation observed in bAM during the first 24 h of *M. bovis* infection ^23^ is due to reconfiguration of CpG methylation patterns. On the other hand, the results presented here indicate that cell signalling and transcription factor-driven gene regulatory transduction cascades lead to the rapid transcriptional activation of immune- and other genes. This observation is supported by previous work showing that DNA methylation changes in THP-1 macrophages infected with *M. tuberculosis* do not occur at CpGs ^14^. In their study, Sharma and colleagues demonstrated that methylation was perturbed at non-CpGs. Similarly, Lyu *et al.* recently demonstrated that infection of human THP-1 macrophages with virulent and avirulent *M. tuberculosis* is not associated with host DNA methylation changes ^45^. Unlike THP-1 macrophages infected with *M. tuberculosis*, we show that non-CpG methylation is not altered in bAM infected with *M. bovis*. The lack of differences in non-CpG methylation may be explained, in part, by the differences in cell types used—comparing a macrophage-like human cell line (THP-1) to a primary, differentiated bovine macrophage. Mycobacteria have recently been reported to modulate the host immune response through chromatin modifications ^46,47^. Given the absence of differential CpG methylation between non-infected and *M. bovis*-infected bAM at 24 hpi, and the differential expression of genes encoding chromatin modifiers observed in the current study, it is reasonable to hypothesise that chromatin reconfiguration may have a role in regulating host gene expression in response to infection with *M. bovis*.

To comprehensively annotate gene promoter methylation in bAM we quantified average DNA methylation at proximal promoter regions spanning the TSS (1,500 bp upstream and 500 bp downstream). As expected, the majority of promoters containing or overlapping a CGI were hypomethylated and those promoters not associated with CGIs were, generally, highly methylated ^48^. However, a large number of promoters (2,580) exhibited mean methylation levels ranging between 33–66% (intermediately methylated; IM). Interestingly, in addition to this, gene ontology analysis of the genes proximal to these promoters indicated a marked enrichment for immune-function related categories. Further analysis of the IM promoter group revealed that most promoters were included due to averaging of methylated and unmethylated CpGs within the 2 kb promoter regions. After removing these promoters, 267 promoters remained that exhibited IM. Validation experiments, using clonal analysis and COBRA, confirmed intermediate DNA methylation at the proximal promoter of two of these non-imprinted IM genes, *C1QB* and *IL2RA*. Six of the 267 promoters were proximal to known bovine imprinted genes, displaying predominant intermediate methylation of 5’ CGIs; as expected for imprinted genes in an adult somatic cell type ^49^. The remaining promoters are IM non-imprinted genes. Previous work by Weber and co-workers demonstrated that, in somatic cells, the concentration of CpGs within a gene promoter is related to the level of DNA methylation; promoters with a high frequency of CpGs (HCP) tend to be unmethylated and promoters with a lower CpG content (LCP) tend to be methylated ^48^. Sixty of the 267 IM promoters identified in this study contained high frequencies of CpGs (CGIs) normally associated with unmethylated HCPs, suggesting that promoter IM in bAM is occurring irrespective of CpG density. It has been suggested that intermediate DNA methylation is a conserved signature of genome regulation associated with intermediately active rather than suppressed gene expression ^41^. It is possible that these intermediately methylated promoters are a hallmark of bAM and functionally associated with this particular cell type. However, Elliot and colleagues demonstrated that different tissues and cell types are intermediately methylated equally ^41^; therefore, the function of intermediate methylation at these genomic loci remains to be fully elucidated.

## Conclusion

This is the first comprehensive analysis of the mammalian alveolar macrophage DNA methylome in response to infection with a mycobacterial pathogen. Although the epigenome of host bAM was not perturbed by a 24 h exposure to the pathogenic bacterium, *M. bovis*, this work provides the first annotation of genome-wide DNA methylation patterns in the bovine genome and is directly aligned with the goal of the Functional Annotation of Animal Genomes (FAANG) project to ‘*produce comprehensive maps of functional elements in the genomes of domesticated animal species*’ ^50^. Furthermore, this work also provides evidence for differential methylation at the proximal promoter regions of more than 200 non-imprinted genes.

## Acknowledgments

We would like to thank Felix Krueger of the Babraham Institute for advice on WGBS data processing and quality control and for providing permission to use the Bismark Reports and Babraham Bioinformatics logos, located in the supplementary information. In addition, we would like to thank Han Haige for assistance with translation of a Chinese journal article.

## Author Contributions

Conceived and designed the experiments: AMOD, DAM, SVG and DEM. Prepared the samples: DAM and JB. Performed the experiments: AMOD, DAM, SA and REI. Provided RNA-seq data: NCN. Analysed the data: KRA, MWW, AMOD, SA and TJH. Prepared the manuscript: AMOD, KRA and DEM. All authors read and approved the manuscript.

## Data availability

All WGBS data is available from NCBI GEO ^51^ (Accession Number GSE110412).

## References

1 Garnier, T. et al. The complete genome sequence of *Mycobacterium bovis*. Proc. Natl. Acad. Sci. U. S. A. 100, 7877–7882, doi:10.1073/pnas.1130426100 (2003).

2 Olea-Popelka, F. et al. Zoonotic tuberculosis in human beings caused by *Mycobacterium bovis*-a call for action. Lancet Infect. Dis. 17, e21-e25, doi:10.1016/S1473-3099(16)30139-6 (2017).

3 Kathirvel, M. & Mahadevan, S. The role of epigenetics in tuberculosis infection. Epigenomics 8, 537–549, doi:10.2217/epi.16.1 (2016).

4 Holoch, D. & Moazed, D. RNA-mediated epigenetic regulation of gene expression. Nat. Rev. Genet. 16, 71–84, doi:10.1038/nrg3863 (2015).

5 Jaenisch, R. & Bird, A. Epigenetic regulation of gene expression: how the genome integrates intrinsic and environmental signals. Nat. Genet. 33 **Suppl**, 245–254, doi:10.1038/ng1089 (2003).

6 O’Doherty, A. M. & McGettigan, P. A. Epigenetic processes in the male germline. Reprod. Fertil. Dev. 27, 725–738, doi:10.1071/rd14167 (2015).

7 Yong, W. S., Hsu, F. M. & Chen, P. Y. Profiling genome-wide DNA methylation. Epigenetics Chromatin 9, 26, doi:10.1186/s13072-016-0075-3 (2016).

8 Gonzalez, R. M., Ricardi, M. M. & Iusem, N. D. Epigenetic marks in an adaptive water stress-responsive gene in tomato roots under normal and drought conditions. Epigenetics 8, 864–872, doi:10.4161/epi.25524 (2013).

9 Kucharski, R., Maleszka, J., Foret, S. & Maleszka, R. Nutritional control of reproductive status in honeybees via DNA methylation. Science 319, 1827–1830, doi:10.1126/science.1153069 (2008).

10 Navarro-Martin, L. et al. DNA methylation of the gonadal aromatase (*cyp19a*) promoter is involved in temperature-dependent sex ratio shifts in the European sea bass. PLoS Genet. 7, e1002447, doi:10.1371/journal.pgen.1002447 (2011).

11 O’Doherty, A. M. et al. Negative energy balance affects imprint stability in oocytes recovered from postpartum dairy cows. Genomics 104, 177–185, doi:10.1016/j.ygeno.2014.07.006 (2014).

12 Waterland, R. A. et al. Season of conception in rural Gambia affects DNA methylation at putative human metastable epialleles. PLoS Genet. 6, e1001252, doi:10.1371/journal.pgen.1001252 (2010).

13 Esterhuyse, M. M., Linhart, H. G. & Kaufmann, S. H. Can the battle against tuberculosis gain from epigenetic research? Trends Microbiol. 20, 220–226, doi:10.1016/j.tim.2012.03.002 (2012).

14 Sharma, G. et al. Genome-wide non-CpG methylation of the host genome during *M. tuberculosis* infection. Sci. Rep. 6, 25006, doi:10.1038/srep25006 (2016).

15 Zheng, L. et al. Unraveling methylation changes of host macrophages in *Mycobacterium tuberculosis* infection. Tuberculosis (Edinb) 98, 139–148, doi:10.1016/j.tube.2016.03.003 (2016).

16 Shell, S. S. et al. DNA methylation impacts gene expression and ensures hypoxic survival of *Mycobacterium tuberculosis*. PLoS Pathog. 9, e1003419, doi:10.1371/journal.ppat.1003419 (2013).

17 Doherty, R. et al. The CD4(+) T cell methylome contributes to a distinct CD4(+) T cell transcriptional signature in *Mycobacterium bovis*-infected cattle. Sci. Rep. 6, 31014, doi:10.1038/srep31014 (2016).

18 Marr, A. K. et al. *Leishmania donovani* infection causes distinct epigenetic DNA methylation changes in host macrophages. PLoS Pathog. 10, e1004419, doi:10.1371/journal.ppat.1004419 (2014).

19 Sinclair, S. H., Yegnasubramanian, S. & Dumler, J. S. Global DNA methylation changes and differential gene expression in *Anaplasma phagocytophilum*-infected human neutrophils. Clin. Epigenetics 7, 77, doi:10.1186/s13148-015-0105-1 (2015).

20 Sitaraman, R. *Helicobacter pylori* DNA methyltransferases and the epigenetic field effect in cancerization. Front. Microbiol. 5, 115, doi:10.3389/fmicb.2014.00115 (2014).

21 Magee, D. A. et al. Innate cytokine profiling of bovine alveolar macrophages reveals commonalities and divergence in the response to *Mycobacterium bovis* and *Mycobacterium tuberculosis* infection. Tuberculosis (Edinb) 94, 441–450, doi:10.1016/j.tube.2014.04.004 (2014).

22 Malone, K. M. et al. Comparative ‘omics analyses differentiate *Mycobacterium tuberculosis* and *Mycobacterium bovis* and reveal distinct macrophage responses to infection with the human and bovine tubercle bacilli. Microb. Genom., doi:10.1099/mgen.0.000163 (2018).

23 Nalpas, N. C. et al. RNA sequencing provides exquisite insight into the manipulation of the alveolar macrophage by tubercle bacilli. Sci. Rep. 5, 13629, doi:10.1038/srep13629 (2015).

24 Vegh, P. et al. MicroRNA profiling of the bovine alveolar macrophage response to *Mycobacterium bovis* infection suggests pathogen survival is enhanced by microRNA regulation of endocytosis and lysosome trafficking. Tuberculosis (Edinb) 95, 60–67 (2015).

25 O’Doherty, A. M. et al. DNA methylation dynamics at imprinted genes during bovine pre-implantation embryo development. BMC Dev. Biol. 15, 13, doi:10.1186/s12861-015-0060-2 (2015).

26 O’Doherty, A. M. et al. DNA methylation plays an important role in promoter choice and protein production at the mouse *Dnmt3L* locus. Dev. Biol. 356, 411–420, doi:10.1016/j.ydbio.2011.05.665 (2011).

27 O’Doherty, A. M., O’Shea, L. C. & Fair, T. Bovine DNA methylation imprints are established in an oocyte size-specific manner, which are coordinated with the expression of the DNMT3 family proteins. Biol. Reprod. 86, 67, doi:10.1095/biolreprod.111.094946 (2012).

28 Krueger, F. & Andrews, S. R. Bismark: a flexible aligner and methylation caller for Bisulfite-Seq applications. Bioinformatics 27, 1571–1572, doi:10.1093/bioinformatics/btr167 (2011).

29 Langmead, B. & Salzberg, S. L. Fast gapped-read alignment with Bowtie 2. Nat. Methods 9, 357–359, doi:10.1038/nmeth.1923 (2012).

30 Zimin, A. V. et al. A whole-genome assembly of the domestic cow, Bos taurus. Genome Biol. 10, R42, doi:10.1186/gb-2009-10-4-r42 (2009).

31 Hansen, K. D., Langmead, B. & Irizarry, R. A. BSmooth: from whole genome bisulfite sequencing reads to differentially methylated regions. Genome Biol. 13, R83, doi:10.1186/gb-2012-13-10-r83 (2012).

32 Karolchik, D. et al. The UCSC Table Browser data retrieval tool. Nucleic Acids Res. 32, D493-496, doi:10.1093/nar/gkh103 (2004).

33 Akalin, A. et al. methylKit: a comprehensive R package for the analysis of genome-wide DNA methylation profiles. Genome Biol. 13, R87, doi:10.1186/gb-2012-13-10-r87 (2012).

34 Benjamini, Y. & Hochberg, Y. Controlling the false discovery rate – a practical and powerful approach to multiple testing. J. R. Stat. Soc. Ser. B Method. 57, 289–300 (1995).

35 Young, M. D., Wakefield, M. J., Smyth, G. K. & Oshlack, A. Gene ontology analysis for RNA-seq: accounting for selection bias. Genome Biol. 11, R14, doi:10.1186/gb-2010-11-2-r14 (2010).

36 Ziller, M. J., Hansen, K. D., Meissner, A. & Aryee, M. J. Coverage recommendations for methylation analysis by whole-genome bisulfite sequencing. Nat. Methods 12, 230–232, doi:10.1038/nmeth.3152 (2015).

37 Clark, S. J. et al. Genome-wide base-resolution mapping of DNA methylation in single cells using single-cell bisulfite sequencing (scBS-seq). Nat. Protoc. 12, 534–547, doi:10.1038/nprot.2016.187 (2017).

38 Peat, J. R. et al. Genome-wide bisulfite sequencing in zygotes identifies demethylation targets and maps the contribution of TET3 oxidation. Cell Rep. 9, 1990–2000, doi:10.1016/j.celrep.2014.11.034 (2014).

39 Chen, Z. X. & Riggs, A. D. DNA methylation and demethylation in mammals. J. Biol. Chem. 286, 18347–18353, doi:10.1074/jbc.R110.205286 (2011).

40 Deaton, A. M. et al. Cell type-specific DNA methylation at intragenic CpG islands in the immune system. Genome Res. 21, 1074–1086, doi:10.1101/gr.118703.110 (2011).

41 Elliott, G. et al. Intermediate DNA methylation is a conserved signature of genome regulation. Nat. Commun. 6, 6363, doi:10.1038/ncomms7363 (2015).

42 Nestorov, P., Hotz, H. R., Liu, Z. & Peters, A. H. Dynamic expression of chromatin modifiers during developmental transitions in mouse preimplantation embryos. Sci. Rep. 5, 14347, doi:10.1038/srep14347 (2015).

43 Smith, Z. D. & Meissner, A. DNA methylation: roles in mammalian development. Nat. Rev. Genet. 14, 204–220, doi:10.1038/nrg3354 (2013).

44 Jones, P. A. Functions of DNA methylation: islands, start sites, gene bodies and beyond. Nat. Rev. Genet. 13, 484–492, doi:10.1038/nrg3230 (2012).

45 Lyu, L. N., Jia, H. Y., Li, Z. H., Liu, Z. Q. & Zhang, Z. D. [Changes and differences of DNA methylation in human macrophages infected with virulent and avirulent *Mycobacterium tuberculosis*]. Zhonghua Jie He He Hu Xi Za Zhi 40, 509–514, doi:10.3760/cma.j.issn.1001-0939.2017.07.006 (2017).

46 Sharma, G., Upadhyay, S., Srilalitha, M., Nandicoori, V. K. & Khosla, S. The interaction of mycobacterial protein Rv2966c with host chromatin is mediated through non-CpG methylation and histone H3/H4 binding. Nucleic Acids Res. 43, 3922–3937, doi:10.1093/nar/gkv261 (2015).

47 Yaseen, I., Kaur, P., Nandicoori, V. K. & Khosla, S. Mycobacteria modulate host epigenetic machinery by Rv1988 methylation of a non-tail arginine of histone H3. Nat. Commun. 6, 8922, doi:10.1038/ncomms9922 (2015).

48 Weber, M. et al. Distribution, silencing potential and evolutionary impact of promoter DNA methylation in the human genome. Nat. Genet. 39, 457–466, doi:10.1038/ng1990 (2007).

49 Pervjakova, N. et al. Imprinted genes and imprinting control regions show predominant intermediate methylation in adult somatic tissues. Epigenomics 8, 789–799, doi:10.2217/epi.16.8 (2016).

50 Andersson, L. et al. Coordinated international action to accelerate genome-to-phenome with FAANG, the Functional Annotation of Animal Genomes project. Genome Biol. 16, 57, doi:10.1186/s13059-015-0622-4 (2015).

51 Edgar, R., Domrachev, M. & Lash, A. E. Gene Expression Omnibus: NCBI gene expression and hybridization array data repository. Nucleic Acids Res. 30, 207–210 (2002).

